# The nuclear interactome of DYRK1A reveals a functional role in DNA damage repair

**DOI:** 10.1101/432757

**Authors:** Steven E. Guard, Zachary C. Poss, Christopher C. Ebmeier, Maria Pagratis, Dylan J. Taatjes, William M. Old

**Affiliations:** Department of Molecular, Cellular and Developmental Biology, University of Colorado, Boulder, CO; Department of Biochemistry, University of Colorado, Boulder, CO; Linda Crnic Institute for Down Syndrome, University of Colorado School of Medicine, Anschutz Medical Campus, Aurora, CO

**Author notes:** Corresponding Author: William M. Old, Molecular, Cellular and Developmental Biology, University of Colorado Boulder, Boulder, CO 80309; E.

## Abstract

Loss of function mutations in the protein kinase DYRK1A lead to a syndromic form of autism spectrum disorder and intellectual disability. Conversely, increased DYRK1A dosage is implicated in atypical brain development and neurocognitive deficits in trisomy 21. DYRK1A regulates a diverse array of cellular processes through kinase dependent and independent interactions with substrates and binding partners. Recent evidence implicates DYRK1A in direct regulation of the transcriptional machinery, but many of the molecular details are not yet known. Furthermore, the landscape of DYRK1A interactions in the nucleus is incomplete, impeding progress toward understanding its function in transcription. Here, we used immunoaffinity purification and mass spectrometry to identify nuclear interaction partners of endogenous DYRK1A. These were enriched in DNA damage repair factors, transcriptional elongation factors and E3 ubiquitin ligases. We validated an interaction with RNF169, a factor that promotes homology directed repair upon DNA damage. We further show that knockout of DYRK1A or treatment with DYRK1A inhibitors in HeLa cells impaired efficient recruitment of 53BP1 to DNA double strand breaks induced by ionizing radiation. This nuclear interactome thus reveals a new role for DYRK1A in DNA damage repair and provides a resource for exploring new functions of DYRK1A in the nucleus.

## Introduction

The chromosome 21 encoded protein kinase DYRK1A is essential for brain development in flies, rodents and humans. De novo *DYRK1A* mutations cause intellectual disability, neurodevelopmental defects, and a syndromic form of autism^1,2^. Overexpression of the protein kinase in the developing brain is implicated in the neurocognitive deficits in individuals with trisomy 21(Down syndrome)^3,4^. Normalization of DYRK1A expression or activity in trisomy 21 mouse models have demonstrated rescue of neurodevelopmental and cognitive deficits^5–8^. DYRK1A is pleiotropic, with functions and interactions that depend on subcellular localization, cell type and expression levels^9,10^. DYRK1A kinase activity is thought to be constitutive once activated by co-translational autophosphorylation of an activation-loop tyrosine^11,12^, and is regulated instead through compartmentalization^13^, transcriptional control^14^, and protein-protein interactions^15^. Kinase-independent scaffold mechanisms have been reported for DYRK1A in transcriptional regulation^16,17^, and for DYRK2 as an ubiquitin ligase adapter and substrate kinase^18^. Detailed investigation into the complexes and interactions of scaffolding kinases has yielded important insights into their biological functions^19,20^.

While DYRK1A has established cytosolic roles in regulating the cell cycle and cytoskeleton, its functions within the nucleus are more enigmatic. DYRK1A contains a bipartite nuclear localization signal (NLS) within its kinase domain that is required for nuclear localization, and a C-terminal poly-histidine tract that is required for localization to nuclear speckles^21^. Within these phase-separated nuclear speckle compartments, phosphorylation of various SRSF splicing factors by DYRK1A has been shown to regulate alternative splicing of Tau^22^. DYRK1A was later reported to regulate transcription machinery through kinase dependent and independent interactions with RNA polymerase (Pol) II C-terminal domain (CTD)^23,24^. Despite the accumulating reports of new functions for DYRK1A in the nucleus, many of the molecular interactions underlying these functions remain unknown.

Most DYRK1A protein interactions reported to date were discovered in large-scale interactome studies by affinity-purification of ectopically expressed fusion constructs, followed by mass spectrometry analysis^19,20,25^. Affinity purification mass spectrometry (AP-MS) has enabled large-scale interrogation of the human protein-protein interactome, providing insights into function for the large fraction of the proteome that has no functional annotation^26^. AP-MS studies commonly employ ectopic expression systems that lack the regulatory elements and local chromatin environment of the gene of interest. Consequently, consistent expression levels are difficult to maintain near endogenous levels, which may disrupt stoichiometric balances for multiprotein complexes and pathways, particularly for dosage-sensitive genes^27–29^. Non-physiological overexpression of DYRK1A has been shown to alter its subcellular distribution^30^, confounding the interpretation of DYRK1A interaction studies that employ ectopic expression systems.

To mitigate these issues, we performed mass spectrometry analysis of immunoaffinity-purified, endogenous DYRK1A from HeLa nuclear extracts. The resulting interactome revealed many previously unreported interactions, representing a significant increase in the number of known DYRK1A interactors. We identified central regulators of DNA damage repair and transcription, including RNF169, members of the BRCA1-A complex, and four subunits of the super elongation complex, consistent with emerging evidence for DYRK1A-dependent regulation of these processes^23,31^. We further show that knockout of DYRK1A or treatment with DYRK1A inhibitors delayed resolution of double stranded breaks after ionizing radiation induced damage in HeLa cells.

## RESULTS

### Nuclear interactome of endogenous DYRK1A

To identify interaction partners of endogenous, nuclear-localized DYRK1A, we immuno-purified DYRK1A from a large-scale preparation of HeLa cell nuclear extracts using four different antibodies in triplicate, followed by label-free mass spectrometry analysis (IP-MS) (Fig. 1A). The four commercial DYRK1A antibodies recognize different epitopes in the N-terminal and C-terminal regions of human DYRK1A (Fig. 1B; Supplementary Table S1). This strategy ensures maximal coverage of interaction partners in the event that the antigenic surface overlaps with a protein interaction interface and disrupts capture of endogenous prey interactions. Immunoprecipitated proteins were digested into tryptic peptides using filter aided sample prep (FASP)^32^ and analyzed by 1D liquid chromatography tandem mass spectrometry (LC/MS/MS) using Orbitrap Fusion instrumentation.

**Figure 1.**
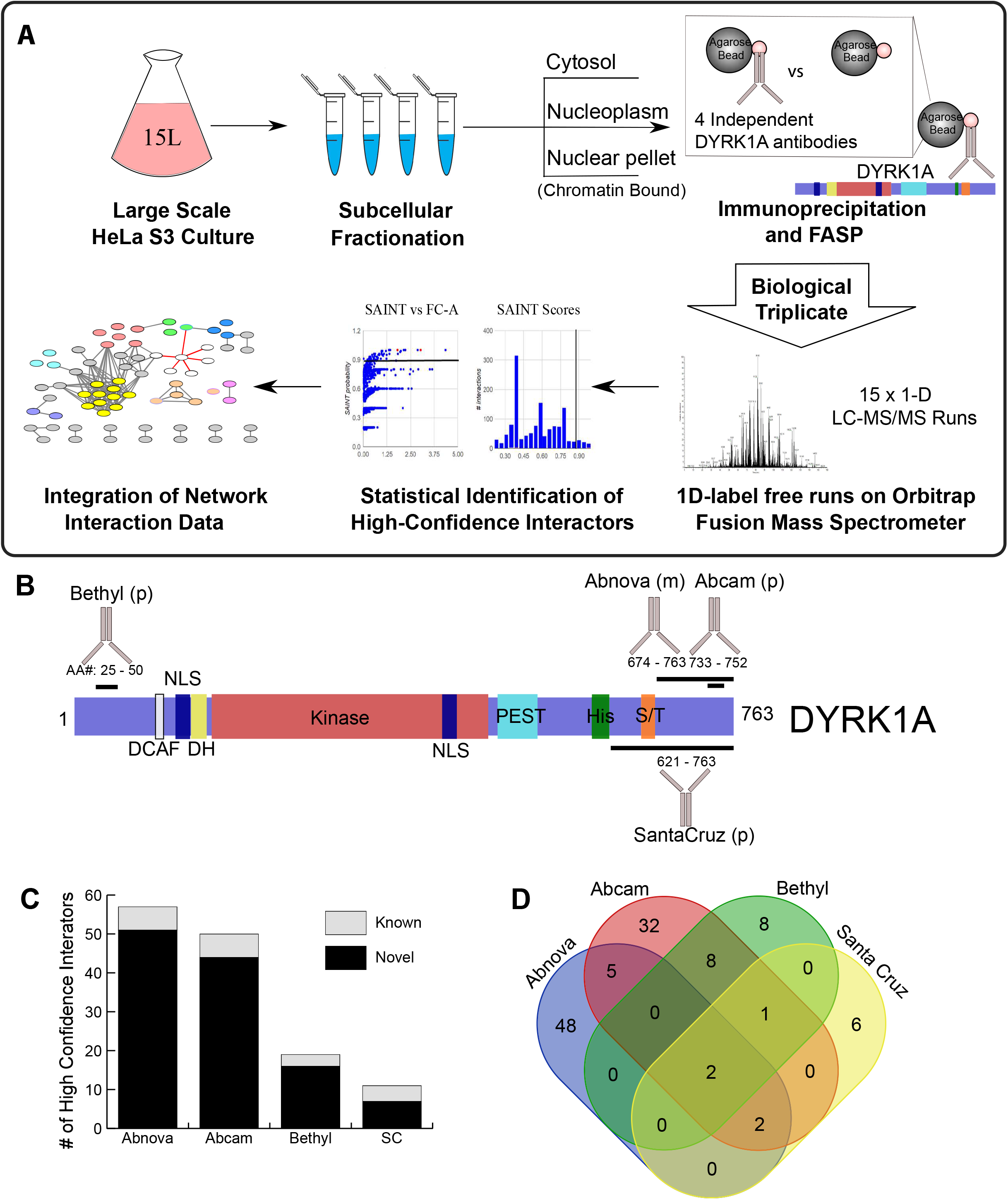
Workflow and strategy for generation of endogenous DYRK1A interactome. **(A)** Schematic overview of fractionation and IPMS strategy. HeLa S3 cells were grown in 6L flasks before harvesting at 1 million cells/ ml. Large scale fractionation of cell pellets generated protein lysates representing a cytosolic, nuclear and chromatin bound fractions. Subsequent immunoprecipitations were done in triplicate prior to sample preparation for mass spectrometry identification. **(B)** Domain arrangement of DYRK1A and regions containing antibody epitope as reported by manufacturers. NLS, nuclear localization signal; DH, DYRK homology box; PEST, proline glutamic acid serine and threonine rich sequence; His, polyhistidine stretch; S/T serine and threonine rich region. Epitopes recognized by each antibody used for IP experiments fall within the designated amino acid region and represented by black bar. (p) rabbit polyclonal; (m) mouse monoclonal. **(C)** Number of novel and literature reported HCI’s by antibody. (Abnova n = 3; Abcam n = 3; Bethyl n = 3; SC: Santa Cruz n = 3) FC-A > 3; SAINT > 0.7. **(D)** Overlap between HCI’s of each set of immunoprecipitations by antibody.

In affinity proteomics experiments, quantification of bait-prey interactions relative to control is critical for distinguishing true interacting proteins from non-specific background. This is particularly important in affinity purifications from complex lysates, in which non-specific interactions predominate as a function of total protein^33^. To distinguish true DYRK1A interacting proteins from non-specific background, we used beads only controls and the CRAPome analysis tool^33^. This approach calculates two measures: a posterior probability of true interaction using the SAINT algorithm^34^, and a fold-change enrichment, FC-A, which estimates an enrichment over internal user controls. High confidence interactions (HCIs) were defined as proteins with an FC-A of 3.0 or greater and a SAINT probability of 0.7 or greater. Our analysis revealed a total of 105 HCIs (Fig. 1C,D; Supplementary Table S2), 97 of which have not been reported as DYRK1A interacting proteins (Fig. 1C). For each antibody, DYRK1A was identified within the four most highly enriched proteins (ranked by FC-A), indicating high-specificity toward DYRK1A from nuclear extracts. A core set of 5 HCIs were shared by three or more antibodies (Fig. 1D), which included the known DYRK1A interacting proteins DCAF7, GLCCI1, RNF169, TROAP, and FAM117B^20,35^. While the Abnova and Abcam antibodies resulted in the identification of several-fold more interactions than the SantaCruz and Bethyl antibodies, most represented novel interactions (Fig. 1C). Many HCIs were unique to the Abnova and Abcam antibodies (Fig. 1D), suggesting that these antibodies recognize distinct DYRK1A sub-complexes, potentially due to epitope overlap within a protein-protein interaction surface.

To gain insight into biological functions associated with the DYRK1A HCIs, we performed functional enrichment and STRING network analysis. HCIs identified in each of the four sets of immunoprecipitations were combined and mapped onto a STRING protein-protein interaction network (Fig. 2A). Functional enrichment analysis using ClueGO^36^ revealed several distinct clusters of functionally related proteins and multiprotein complexes involved in transcription and DNA damage repair (Fig. 2B). For example, we identified three of the five subunits of the BRCA1-A complex, which plays key roles in the repair of DNA damage induced by ionizing radiation^37,38^, as well as TRIP12, RAD18 and ERCC5 (Fig. 2A). RAD18 is a RING-type E3 ligase that has been reported to promote HR and antagonize NHEJ^39^. TRIP12 is a HECT-type E3 ubiquitin ligase that limits spreading of ubiquitylation marks surrounding DSBs by suppressing the accumulation of RNF168 in a UBR5 dependent manner^40^. Interestingly, haploinsufficiency of TRIP12 has been reported to cause intellectual disability^41^.

**Figure 2:**
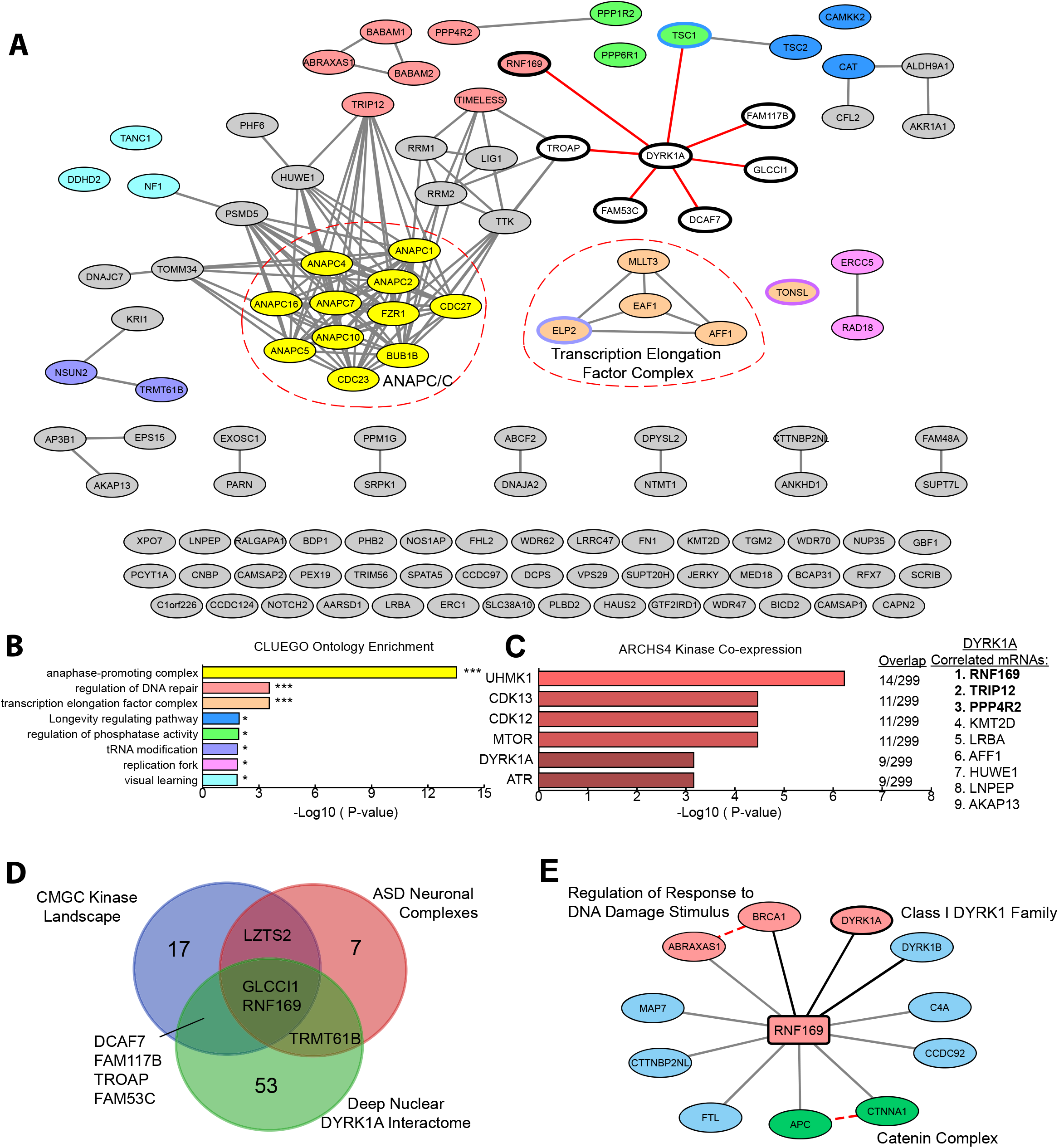
Network analysis of protein-protein interactions (PPIs) from nuclear DYRK1A interactome. **(A)** STRING-DB network of high significance PPIs from nuclear DYRK1A interactome. Nodes in network represent proteins found in a shared HCI list between antibodies. Total Nodes: 106. HCI cutoffs: FC-A > 3, SAINT > 0.7. Black Edges denote STRING-DB evidence of 0.400 for interactions between HCIs. Red edges denote confirmed BioGRID interactions. Color of node correlates to GO ontologies outlined in figure 2b. **(B)** ClueGO ontology enrichments from total HeLa nuclear DYRK1A interactome. (Fisher exact test: * p < 0.05; ** p < 0.01; *** p < 0.001). **(C)** Enrichment of HCIs against the ARCHS4 RNA-seq database for kinase co-expression. (Lists of top 300 most correlated mRNAs for each kinase are generated through Enrichr.) Bold: GO:2000779 Regulation of double-strand break repair (Fisher exact test: adj. p = 0.00004). **(D)** Comparison of DYRK1A interactomes with literature reported interaction sets. Green: Nuclear DYRK1A HCI interactome; Blue: Varjosalo *et al*.^20^; Red: Li *et al*. (2015)^49^. **(E)** RNF169 nuclear interactome (FC-A>3; SAINT > 0.7) n = 2. Grey edges denote high confidence interactions; Red edges: STRING-DB evidence 0.400. Regulation of response to DNA damage stimulus p = 0.01; Catenin complex p = 0.007.

The most significantly enriched term from the ClueGO analysis was related to the anaphase-promoting complex (APC/C) (Fig. 2B). We identified 10 of the 15 core subunits of APC/C (Fig. 2A,B, Suppl. Fig. S1A) with significant enrichment over the beads-only control IPs. APC/C is a 1.2 MDa E3 ubiquitin ligase complex that controls mitotic progression by regulating the activity of cyclin-dependent kinases^42^ and has been recently implicated in regulation of DNA damage repair pathway choice^43^. Substrate specificity of APC/C is determined by mutually exclusive association with the adaptor proteins FZR1 (Cdh1) and CDC20, which bind APC/C in a cell cycle dependent manner^44^. Interestingly, in our list of DYRK1A HCIs, we found FZR1 (Cdh1 homolog 1), but not CDC20, suggesting that DYRK1A preferentially interacts with the G1-specific APC/C-Cdh1 complex^45^.

The mRNA expression levels for physically interacting proteins have been shown to co-evolve, and as a result show high pairwise expression correlation^46^. Given a suitably large compendium of expression data, these correlation patterns can be detected using various co-expression analysis techniques, and then used to infer protein-protein interactions. This has been demonstrated by large-scale correlation analysis of the entire Gene Expression Omnibus collection of RNA-seq experiments across thousands of human cell types and perturbations, which showed high accuracy prediction of protein interactions and KEGG pathway membership^47^.

Using this principle, we asked whether enrichment of co-regulated mRNAs in our interactome might reveal stable interactions within our DYRK1A HCIs. We tested for significant overlap of proteins in our DYRK1A interactome with ARCHS4 kinase co-expression modules using Enrichr^48^. We found that DYRK1A HCIs were enriched in mRNAs that show high expression correlation with the cyclin dependent kinases CDK12 and CDK13, as well as ATR, which acts as a DNA damage sensor to initiate recruitment of repair proteins. As expected, we found that the module of DYRK1A co-regulated mRNAs showed significant overlap with DYRK1A HCIs. Examination of these mRNAs revealed that many were involved in DNA damage (Fig. 2C). This analysis suggests that DYRK1A may be involved in DNA damage repair, potentially through association with complexes that are recruited to DNA double stranded breaks (DSBs).

### Interaction with RNF169 implicates DYRK1A in DNA damage repair

RNF169 was consistently represented with the highest combination of SAINT scores and FC-A values across every replicate of all four DYRK1A antibodies used in this study (Suppl. Fig. S1B-E). This interaction was previously reported in two independent proteomic studies using HEK293T and SH-SY5Y cells (Fig. 2D)^20,49^. RNF169 has emerged as an important factor influencing DNA repair pathway outcome upon DSB formation, and has been shown to promote high-fidelity homologous recombination (HR) repair^50,51^ and single-strand annealing (SSA) repair^52^. 53BP1 recruitment to histone H2A ubiquitin marks placed by the RNF169 paralog, RNF168, inhibits DNA end resection and promotes non-homologous end joining (NHEJ)^53,54^. RNF169 and the BRCA1-RAP80 complex compete with 53BP1 for binding to these H2A ubiquitin marks, thereby antagonizing NHEJ in favor of HR and SSA repair pathways^52,55,56^.

To validate the DYRK1A-RNF169 interaction and identify proteins that may interact with RNF169, we performed IP-MS on RNF169 from HeLa nuclear extracts. Consistent with the DYRK1A IP-MS data, DYRK1A was identified in RNF169 IPs, in addition to DYRK1B, a class I DYRK kinase. Interestingly, DYRK1B was not found in the DYRK1A IP-MS data, suggesting mutually exclusive binding of DYRK1A and DYRK1B to a common interaction surface of RNF169. Among the RNF169 HCIs were subunits of the BRCA1-A complex: ABRAXAS1 and BRCA1 (Fig. 2E). This complex was well represented in the DYRK1A interactome by proteins ABRAXAS1, BABAM1 (also known as NBA1 and MERIT40) and BABAM2 (also known as BRE and BRCC45) (Fig. 2A). The shared interaction of both DYRK1A and RNF169 with BRCA1-A subunits is interesting in light of the requirement for this complex in efficient homology directed repair of DSBs^37,57–59^.

### DYRK1A levels influence DNA DSB repair protein recruitment

The DYRK1A nuclear interactome was enriched in DSB repair regulators, particularly factors involved in antagonizing NHEJ, suggesting that DYRK1A could play a role in regulating NHEJ efficiency. To test this idea, we examined the effect of DYRK1A knockout, or treatment with DYRK1A inhibitors, on the recruitment and resolution of 53BP1 to sites of DNA damage induced by ionizing radiation (IR) in HeLa cells (Fig. 3A). 53BP1 is a driver of non-homologous end joining and is recruited to double-strand breaks within 15 minutes of IR induced damage^56^. We tested three structurally unrelated DYRK1A inhibitors (harmine, L41, and INDY) across a range of doses spanning their reported EC50s. We also generated a DYRK1A KO HeLa cell line by CRIPSR/Cas9 gene editing (Fig. 3B). WT and DYRK1A KO cells were pre-treated with kinase inhibitor and irradiated at 4 Gy, a dose selected to induce sufficient damage while minimizing widespread apoptosis. Following induction of double strand breaks by IR, cells were fixed at 1, 4 and 8 hours and stained for γH2AX, 53BP1 and Hoechst to visualize and quantify the formation and resolution of IR induced foci.

**Figure 3:**
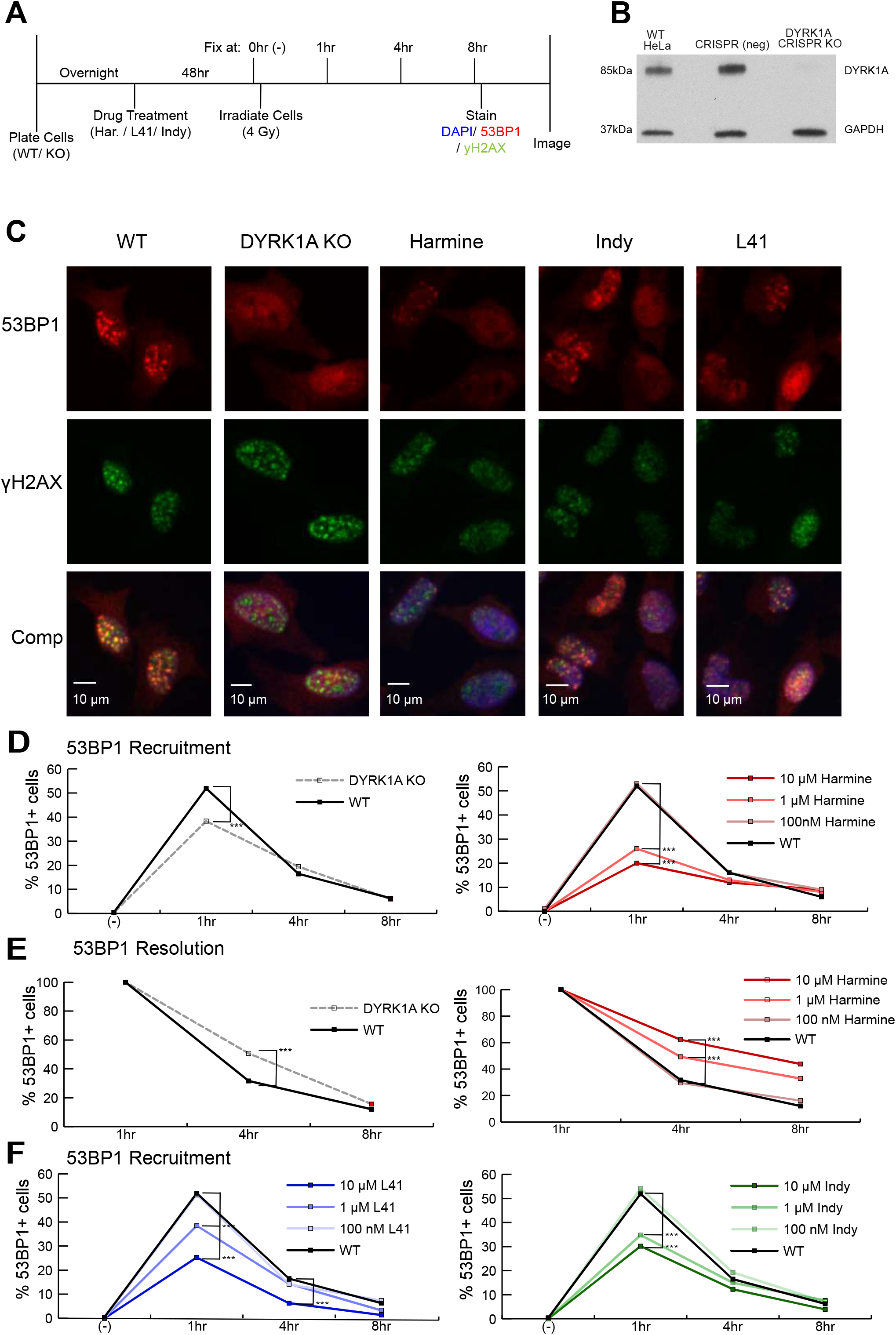
DYRK1A expression and kinase activity are necessary for efficient 53BP1 recruitment to double-strand break sites. **(A)** Experimental timeline for 53BP1 foci quantitation in response to DYRK1A manipulation and ionizing radiation. Cells were treated with drug 48 hours prior to treatment with ionizing radiation (4 Gy). Cells were fixed immediately before (0 hr) or 1, 4 and 8 hours following ionizing radiation. **(B)** Western blot confirmation of DYRK1A CRISPR knockout in HeLa cells. GAPDH loading control. **(C)** Representative immunofluorescent images of yH2AX and 53BP1 staining in fixed HeLa cells. Cells were fixed in 96-well plates. Four frames per well were imaged for each well. Four wells per condition per time point were plated and quantified using the Focinator R package^77^. N ≈ 400 – 1000 cells per condition. Blue: Hoechst; Green: yH2AX; Red: 53BP1. yH2AX+ cells ≥ 10 foci/ cell; 53BP1+ cells ≥ 10 foci/ cell. (53BP1 noise cut off: 15; yH2AX noise cut off 20). **(D)** DYRK1A KO and harmine treated HeLa cells: Proportion of 53BP1+ cells over time following 4 Gy of ionizing radiation. (KO: DYRK1A HeLa CRISPR KO cells.) **(E)** Resolution of 53BP1 foci in DYRK1A KO cells and harmine treated HeLa cells vs WT: The number of 53BP1+ cells were normalized to the maximal recruitment at 1 hr. Resolution of these foci were measured ratiometrically between the 53BP1+ cells at 1 hr:4 hr and 1 hr:8 hr. (Proportion test; *: p <0.05; **: p<0.01; ***: p<0.001). **(F)** L41 and INDY treated HeLa cells: Proportion of 53BP1+ cells over time following 4 Gy of ionizing radiation.

We found that in HeLa cells, 53BP1 recruitment maximized one-hour post IR (Suppl. Fig. S2A). Resolution of 53BP1 foci was measured by quantifying the decrease in foci per cell at four-hour and eight-hour time points, compared to the maximum recruitment levels. We observed that unirradiated HeLa cells typically contain basal levels of γH2AX foci and singular, large 53BP1 foci characteristic of stalled replication forks and inherited DNA damage lesions (Suppl. Fig. S2A)^60^. To quantify the number of cells with ionizing radiation induced foci, we defined 53BP1 positive cells as containing greater than 10 foci per nucleus, as described previously^51^.

Relative to the parental HeLa line, DYRK1A KO cells exhibited a decreased proportion of 53BP1+ cells at one hour post IR, consistent with impaired recruitment of 53BP1 to ionizing radiation induced foci (Fig. 3C,D). Further, DYRK1A knockout cells showed an increased proportion of 53BP1+ cells at 4 hours post IR (normalized to 1 hr), relative to the parental line, indicating a decreased rate in resolution of 53BP1 foci (Fig. 3E). Similar to knockout of DYRK1A, pre-treatment with the DYRK1A inhibitor, harmine, led to a significant, dose-dependent reduction of 53BP1 recruitment for both recruitment and resolution (Fig. 3D,E). Treatment with the structurally unrelated DYRK1A inhibitors, L41 and INDY, phenocopied both the harmine treatment and DYRK1A KO in 53BP1 recruitment, albeit to different extents. (Fig. 3F). All three drugs inhibit DYRK1A in an ATP competitive manner but have unique off-target profiles^61,62^. In contrast to harmine treated cells and DYRK1A knockout cells, the rate of resolution for 53BP1 foci in L41 and INDY treated cells did not significantly differ from control cells (Suppl. Fig. 2B,C), suggesting that DYRK1A is more likely involved in 53BP1 recruitment than resolution. As removal of 53BP1 by HR repair proteins is cell cycle dependent, resolution rates of DNA double strand breaks at later time points could confounded by DYRK1A dependent shifts in cell cycle^63^. Consistent with prior studies, quantification of cell cycle phase by flow cytometry showed that the fraction of cells in S-phase was increased slightly in DYRK1A KO cells relative to the control line (Suppl. Fig. S3D)^64^. Taken together, these results indicate that DYRK1A expression and activity are required for efficient 53BP1 recruitment to DSBs within one hour, which contrasts with the reported role for RNF169 in antagonizing 53BP1 recruitment to promote HR and SSA^50,52^.

### DYRK1A independent regulation of 53BP1 recruitment

To test whether DYRK1A mediates the harmine-induced 53BP1 phenotype in response to IR, we treated DYRK1A KO cells with harmine and measured 53BP1 recruitment in response to IR. Surprisingly, 53BP1 recruitment was further reduced in a dose dependent manner (Fig. 4A). The closely related kinase, DYRK1B, is inhibited by INDY and L41 at approximately the same concentration as DYRK1A, and is inhibited by harmine with an IC50 2–3 times higher than that for DYRK1A^61,62,65,66^. As both DYRK1A and DYRK1B were identified as RNF169 HCIs (Fig. 2E), we speculate that the catalytic activity of these kinases could be acting in a semi-redundant manner to regulate RNF169.

**Figure 4:**
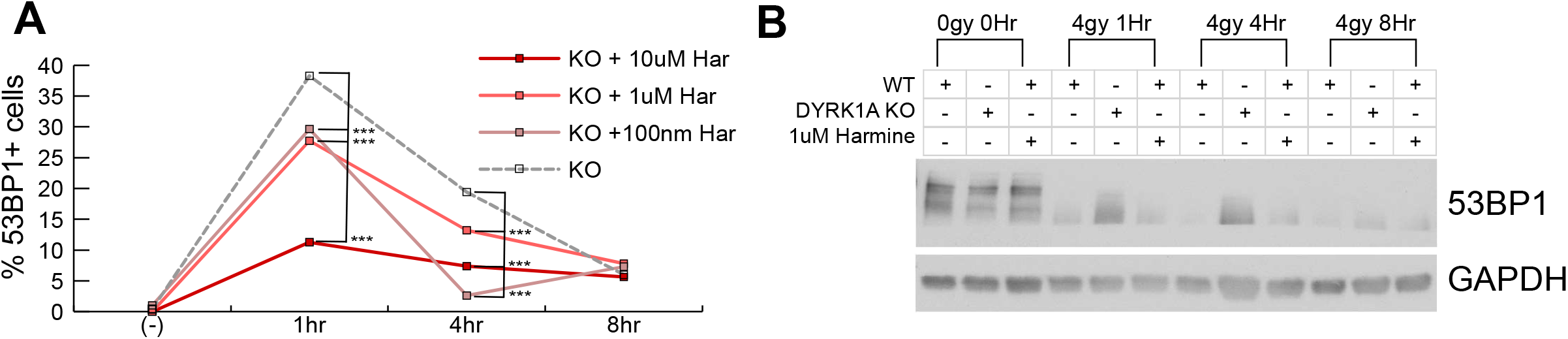
Class 1 DYRK inhibitors exacerbate 53BP1 recruitment phenotype in DYRK1A KO cells. **(A)**.DYRK1A KO cells were treated with 100 nM, 1 µM or 10 µM harmine 48 hours prior to ionizing radiation. Cells were fixed and quantified as in Fig. 3A. (Proportion test: drug treated cells vs DYRK1A KO; *: p <0.05; **: p<0.01; ***: p<0.001). **(B)** Immunoblot for 53BP1 degradation and GAPDH expression in either WT, DYRK1A KO or 1 µM harmine treated WT cells at 0 hrs, 1 hr, 4 hr and 8 hrs following 4 Gy of IR.

Collectively, our data indicate that loss of DYRK1A expression or kinase activity impairs the recruitment of 53BP1 to double strand break sites. Upon IR induced DNA damage, nucleoplasmic 53BP1 is rapidly degraded by the proteasome, while chromatin-localized 53BP1 is protected from degradation^67^. We observed that IR-induced 53BP1 degradation was impaired in the DYRK1A KO line relative to the parental control line, while harmine treated parental control cells showed no effect (Fig. 4B). This points to both kinase dependent and independent mechanisms for DYRK1A in regulation of 53BP1 stability after IR induced DSB formation.

## DISCUSSION

The molecular mechanisms that underlie the dosage sensitivity of DYRK1A during human brain development remain unclear, but likely involve perturbations to DYRK1A interaction networks and phosphorylation targets. The interactors of human DYRK1A described to date have been found by low throughput biochemical methods (see Table S2 in ^68^) or large-scale interactome AP-MS analysis^20,25,49^. Compared with DYRK1A cytosolic interactions, much less is known about its protein interactions in the nucleus, where it is known to localize in human, mouse and chicken neuronal cells^10,69,70^.

In this study, we performed a deep proteomic analysis of DYRK1A-associated proteins in the nucleus by mass spectrometry analysis of endogenous DYRK1A, immunopurified from a large-scale preparation of HeLa cell nuclear extract. We identified 105 high-confidence DYRK1A interactions, only 8 of which have been described previously, underscoring how current knowledge of DYRK1A interactions is incomplete. This issue is not limited to DYRK1A. A recent AP-MS interactome study of all human CMGC kinases found that more than 75% of the discovered interactions were previously unknown^20^. Enrichment of DYRK1A from nuclear extracts in this study likely contributed to the large number of unique interactions compared with previous studies, which used whole cell lysates^20,49^. The nucleus contains approximately 15% of the total cellular protein in HeLa cells^71^, and accordingly, low abundance proteins from this compartment are likely to be under sampled relative to more abundant cytosolic interactions reported in previous AP-MS experiments using whole cell lysates^20,49^. Although nuclear DYRK1A was recently reported to account for 10% of the total DYRK1A in HeLa cells^23^, we found that it was much more concentrated in the nuclear and chromatin bound fractions than in the cytosol (Suppl. Fig. S1H). Further characterization of these interacting partners will be required to distinguish between primary and secondary interactions.

A small set of DYRK1A HCIs identified here have been independently identified by multiple studies: RNF169, FAM117B, DYRK1A, GLCCI1, TSC1, DCAF7, TROAP, FAM53C and TRMT61B (Fig. 2A,D). These represent a core set of conserved, high affinity interaction partners of DYRK1A, due to their consistent appearance across different cell types employed in previous affinity purification studies. Remarkably, four of these interactors, TROAP, GLCCI1, FAM53C and FAM117B, have very limited functional annotation. This highlights how our understanding of human DYRK1A function remains limited, nearly three decades since the discovery of Yak1p, the founding member of the DYRK family in budding yeast^72^.

DYRK1A was recently reported to stimulate transcriptional elongation of specific genes containing a palindromic promoter element by recruitment to promoter regions and phosphorylation of the C-terminal domain of RNA Pol II^23^. The C-terminal histidine-rich domain of DYRK1A has been shown to promote hyperphosphorylation of the Pol II CTD and phase-separation with Pol II CTD in vitro and in cells^24^. Transcription and pre-mRNA splicing involve the coordinated regulation of large multi-protein molecular machines involving hundreds of factors, raising important questions about other DYRK1A interactions important in transcriptional control. While no RNA pol II interactions were identified in this study, we did identify several subunits of the super elongation complex (SEC), which is involved in rapid transcriptional activation of RNA Pol II^18^. The SEC is also involved in a positive feedback loop with the Notch pathway, which reinforces fate commitment of neural stem cells in *Drosophila* neuroblasts^73^. Intriguingly, we identified NOTCH2 in the set of DYRK1A HCIs (Fig. 2A), raising the possibility that DYRK1A may mediate interactions between the SEC and NOTCH at target genes.

We wondered whether any of the antibodies we used might inform on other studies in the literature using the same reagents. Interestingly, all of the SEC subunits identified as DYRK1A interactions were only detected in DYRK1A IPs using the C-terminal recognizing Abcam antibody (Fig. 1B). The same Abcam and Abnova antibodies from this study were used previously in ChIP experiments to uncover DYRK1A’s transcriptional function^23^. Di Vona *et al*. reported using the Abcam antibody that DYRK1A was recruited to chromatin in HeLa cells while the Abnova DYRK1A antibody prevented binding of DYRK1A to DNA^23^. Taken together, this supports the notion that these antibodies may enrich for a distinct subset of DYRK1A interactions due to differential occlusion of a binding interface, and that DYRK1A recruitment to chromatin could be driven by secondary interactions with elongation factors.

Our DYRK1A interactome illuminates new functions for this protein kinase in DNA damage repair. The DNA damage proteins identified as HCIs in this study clearly implicate DYRK1A in regulating aspects of DNA damage repair, most likely in promoting the recruitment of 53BP1 to the sites of damage. We found that DYRK1A associates with RNF169, which promotes homology directed repair mechanisms and antagonizes 53BP1 foci recruitment. We show that DYRK1A expression is necessary for efficient recruitment of 53BP1 to DNA damage foci and is also required for rapid degradation of 53BP1 following ionizing radiation. Interestingly, the DYRK1A inhibitor, harmine, failed to alter IR-induced 53BP1 degradation, suggesting a kinase independent role for DYRK1A in 53BP1 stability. Harmine treatment of DYRK1A knockout cells further suppressed 53BP1 recruitment to ionizing radiation induced foci, suggesting that other off-target kinases inhibited by harmine, such as DYRK1B, may be involved in this mechanism. It will be crucial to dissect the relative contribution of DYRK-family kinases and other off-targets of harmine in DNA damage repair.

In light of the importance of disruptive mutations in *DYRK1A* in neurodevelopmental disorders^74^, detailed investigation into the interaction partners reported here may reveal new mechanistic insights into DYRK1A function in the developing fetal and adult brain, where expression is particularly high and exhibits dynamic expression changes during fetal development^2^. This nuclear interactome points to new roles for DYRK1A in DNA damage repair and transcriptional regulation that may be important in understanding the contribution of increased DYRK1A copy number in neurocognitive deficits in Down syndrome. The use of cerebral organoids, an emerging human neurodevelopmental model derived from human induced pluripotent stem cells^75^, would enable detailed investigation into the functional role for these protein interactions in a cellular and developmental context relevant to disease pathogenesis.

## Materials and Methods

### Antibodies

The following antibodies were purchased commercially: mouse monoclonal antibodies against DYRK1A (Abnova Corporation, Taipei, Taiwan; H00001859-M01) and phospho-Histone H2A.X (Ser139) (Millipore, Burlington, MA, 05–636), rabbit polyclonal antibodies against DYRK1A (Abcam, Cambridge, UK; ab69811; Bethyl Laboratories, Montgomery, TX; A303–801A; and Santa Cruz Biotechnology, Dallas, TX; sc-28899) and 53BP1 (Abcam; ab21083).

### Cell culture

HeLa cells were obtained from ATCC and cultured in DMEM + Glutamax +10% FBS with antibiotic. DYRK1A knockout in a HeLa cell line was established using a CRISPRCas9 gene engineering method. Cells were transfected with an RFP-tagged Cas9 plasmid and two BFP-tagged sgRNA-plasmids (Sanger lentiviral CRISPR vector U6-gRNA: PGK-puro-2AtagBFP, Sigma) using lipofectamine 3000 (Invitrogen, Carlsbad, CA6). Both sgRNA-plasmids contained guides to DYRK1A exon 5 of either 5’ ATGATCGTGTGGAGCAAGAATGG 3’ (plasmid #1) or 5’ TAAAATAATAAAGAACAAGAAGG 3’(plasmid #2). All plasmids were provided by Josh Molishree, manager of the functional genomics facility at Anschutz Medical Center, University of Colorado, Denver, CO. Transfected cells were FACS-sorted and RFP/BFP positive cells were grown as single-cell clones from a 96 well plate. Clones were screened for DYRK1A KO through western blot, T7 assay and sequencing of targeted region.

### Chemicals and Treatments

DNA damage was induced by exposing cells to 4 Gy of x-ray irradiation using a Faxitron Cabinet X-Ray System (Faxitron, Tucson, AZ). Harmine (SantaCruz), L41 (BioVision, San Francisco, CA), and INDY (Tocris, Bristol, UK) stock solutions were prepared at 10mM in DMSO. Drugs were then diluted down to final concentrations of 100 nM, 1 µM and 10 µM accordingly per experiment.

### Preparation of HeLa nuclear extract and nuclear pellet

HeLa nuclear extract was prepared from isolated nuclei from approximately 1 billion HeLa S3 cells, as described (Dignam et al., 1983)^76^. The insoluble pellet from the nuclear extract (i.e. the nuclear pellet) was solubilized with 100 mM HEPES pH 7.9, 2 mM MgCl2, 100 mM KCl, 20% (v/v) glycerol, protease inhibitors (0.25 mM PMSF, 1 mM Sodium metabisulfite, 1 mM Benzamidine, 1 mM DTT), phosphatase inhibitors (1 µM Microcystin LR (Enzo Lifesciences, Farmingdale, NY), 0.1 mM Sodium orthovanadate, 10 mM beta-glycerophosphate, 5 mM sodium fluoride, 1 mM sodium pyrophosphate (all Sigma)), and nucleases Benzonase (200 U/mL) and DNAse I (50 U/mL). The pellet was chopped, dounce homogenized 20 times and mixed overnight with a stir bar at 4 ºC. The extract was cleared by centrifugation at 14,000 × g and aliquoted for storage at –80 ºC.

### Sample preparation for mass spectrometry

Affinity purified samples were precipitated with the addition of 10% (w/v) porcine insulin (Sigma), 0.1% (w/v) sodium deoxycholate, and 20% (w/v) trichloroacetic acid at 4 ºC. Precipitated protein was pelleted and washed two times with –20 ºC acetone and air dried. Samples were prepared for mass spectrometry using a modified version of the FASP method^32^. Samples were solubilized in 4%(w/v) sodium dodecyl sulfate (SDS), 100 mM Tris pH 8.5, 10 mM TCEP, boiled and allowed to reduce for 20 min, followed by alkylation with 25 mM iodoacetamide for 30 minutes in the dark. The reduced and alkylated proteins were then transferred to a 30 kD MWCO Amicon Ultra (Millipore) ultrafiltration device and concentrated, washed three times with 8 M urea, 100 mM Tris pH 8.5, and again three times with 2 M urea, 100 mM Tris pH 8.5. One microgram endoprotease LysC (Wako, Osaka, Japan) was added and incubated for 3 hrs rocking at ambient temperature, followed by 1 µg trypsin (Promega, Madison, WI), rocking overnight at ambient temperature. Tryptic peptides were collected by centrifugation and desalted using Pierce C-18 spin columns (Thermo Fisher Scientific) and stored dry at –80 ºC.

### Mass spectrometry analysis

Liquid Chromatography/ mass spectrometry analysis. Samples were suspended in 3% (v/v) acetonitrile, 0.1% (v/v) trifluoroacetic acid and direct injected onto a 1.7 µm, 130 Å C18, 75 µm × 250 mm M-class column (Waters), with a Waters M-class UPLC. Tryptic peptides were gradient eluted at 300 nL/ minute, from 3% acetonitrile to 20% acetonitrile in 100 minutes into an Orbitrap Fusion mass spectrometer (Thermo Scientific). Precursor mass spectrums (MS1) were acquired at 120,000 resolution (FWHM) from 380–1500 m/z with an AGC target of 2.0E5 and a maximum injection time of 50 ms. Dynamic exclusion was set for 20 seconds with a mass tolerance of +/− 10 ppm. Isolation for MS2 scans was 1.6 Da using the quadrupole, and the most intense ions were sequenced using Top Speed for a 3 second cycle time. All MS2 sequencing was performed using higher energy collision dissociation (HCD) at 35% collision energy and scanned in the linear ion trap. An AGC target of 1.0E4 and 35 second maximum injection time was used. Raw files were searched against the UniProt human database using MaxQuant version 1.6.1.0 with Cysteine Carbamidomethylation as a fixed modification. Methionine oxidation and protein N-terminal acetylation were searched as variable modifications. All peptides and proteins were thresholded at a 1% false discovery rate (FDR).

### Immunofluorescence

Cells were plated in wells of Corning 96 well plates (#3603) the night before drug treatment. Cells were either treated with 100 nM, 1 µM, or 10 µM harmine/L41/INDY for 48 hours before receiving 4 Gy of irradiation. Cells were then rinsed twice with PBS and fixed with 4% formaldehyde for 20 minutes. Following fixation, antibodies for 53BP1 (Abcam, ab21083), γH2AX (Millipore), and Hoescht (Thermo Fisher Scientific) were used for fluorescent detection in fixed cells. The cells were then imaged with the Yokogawa CellVoyager CV1000 confocal Scanner system using a 20x objective. All wells were imaged using a high throughput program in the CV1000 Acquisition software, allowing for 4 images per well to be taken with multiple confocal planes and computational autofocus. MIP files were then utilized for analysis by R-package Focinator v2^77^. In brief, the number of foci were counted within each nucleus, excluding nuclei on edge of each frame. Li thresholding was used, and noise/ cutoff thresholds were set to 25 for both foci channels. Cells with a nucleus containing ten or more 53BP1 foci are considered 53BP1+ cells. This processing was done on 16 frames per condition resulting in cell counts between 400 and 1000.

### Immunoblotting and immunoprecipitation

HeLa cells were lysed with RIPA buffer, sonicated using a Bioruptor Pico (Diagenode) for 10 cycles of 30 seconds on/ 30 seconds off, and centrifuged at 14,000 × g for 15 minutes. A Pierce BCA protein assay kit was used to determine protein concentrations and samples of 20 µg total protein were resolved using polyacrylamide gels and probed with the corresponding antibody for protein of interest. DYRK1A immunoprecipitations were done in triplicate using one of four antibodies: Abnova H00001859-M01, Abcam 69811, Bethyl A303–801A or Santa Cruz sc-12568. Antibodies were bound to bead- protein A/G mixtures overnight prior to affinity purification from protein lysate. Following a 15 min incubation with benzonase, protein lysates were precleared over protein A/G sepharose beads (GE Healthcare, Chicago, IL) mixture with no antibody for 1 hour. Beads were then spun down and cleared lysate was incubated with Antibody-bead mixture for 4 hours at 4°C. Affinity purified proteins were eluted off of beads using 0.1 M Glycine pH 2.75 twice for 30 minutes.

### Cell Cycle analysis

BrdU was added to HeLa cells growing in culture for 60 minutes prior to trypsinizing and collection. Cells were washed with PBS and fixed using high grade ethanol at –20°C. Cells are treated with 2N HCl/ Triton X-100 for 30 minutes, pelleted and resuspended in Borax to neutralize the sample. Cells were stained with a BrdU-FITC (BioLegend, San Diego California) antibody and propidium iodide (Sigma) for two-dimensional flow cytometry separation of DNA content. Cells were then separated into G0/G1, S, and G2/M phase based on gating of the two parameters.

### siRNA transfection

Silencer select validated siRNAs from Thermo Fisher were used in this study including s4399, s4400 and s4401. Transfections were done using lipofectamine 2000 in OptiMEM + Glutamax. Following 6 hours of incubation with siRNA/lipofectamine mixture, DMEM growth media was replenished. Cells were harvested 24 hours following initial transfection for immunoblotting.

## Acknowledgements

We thank Tom Blumenthal and Joaquin Espinosa for helpful discussions. S.G. was supported by a Blumenthal Predoctoral Fellowship, a Sie Postdoctoral Fellowship and by the NIH Molecular Biophysics Training Program. This work was supported by a Grand Challenge grant to W.M.O from the Linda Crnic Institute for Down syndrome, by a DARPA cooperative agreement 13-34-RTA-FP-007 to W.M.O, and by the NIH (GM110064, to DJT),

## Author Contributions

S.G., Z.P., and W.M.O. conceived and designed the project. S.G. designed and performed the experiments. Z.P., C.E. and W.M.O contributed to experimental design. C.E. acquired and analyzed the raw mass spectrometry data, S.G. performed the statistical analysis of mass spectrometry and imaging data. M.P. generated the DYRK1A K.O. HeLa cell line. D.J.T. provided HeLa nuclei for proteomics experiments. S.G. and W.M.O wrote the manuscript. All coauthors contributed to editing and reviewing the manuscript.

## Data availability statement

The datasets generated during and/or analyzed during the current study are available in the MassIVE repository: ftp://massive.ucsd.edu/MSV000082881

## Supplementary Files

**Supplementary Table 1**. Table of antibody registry information for DYRK1A antibodies used in immunoprecipitation experiments

**Supplementary Table 2**. DYRK1A nuclear interactions identified in this study and collected from literature

**Supplementary Table 3**. Table of fluorescent foci founts

**Supplemental Figure 1.**
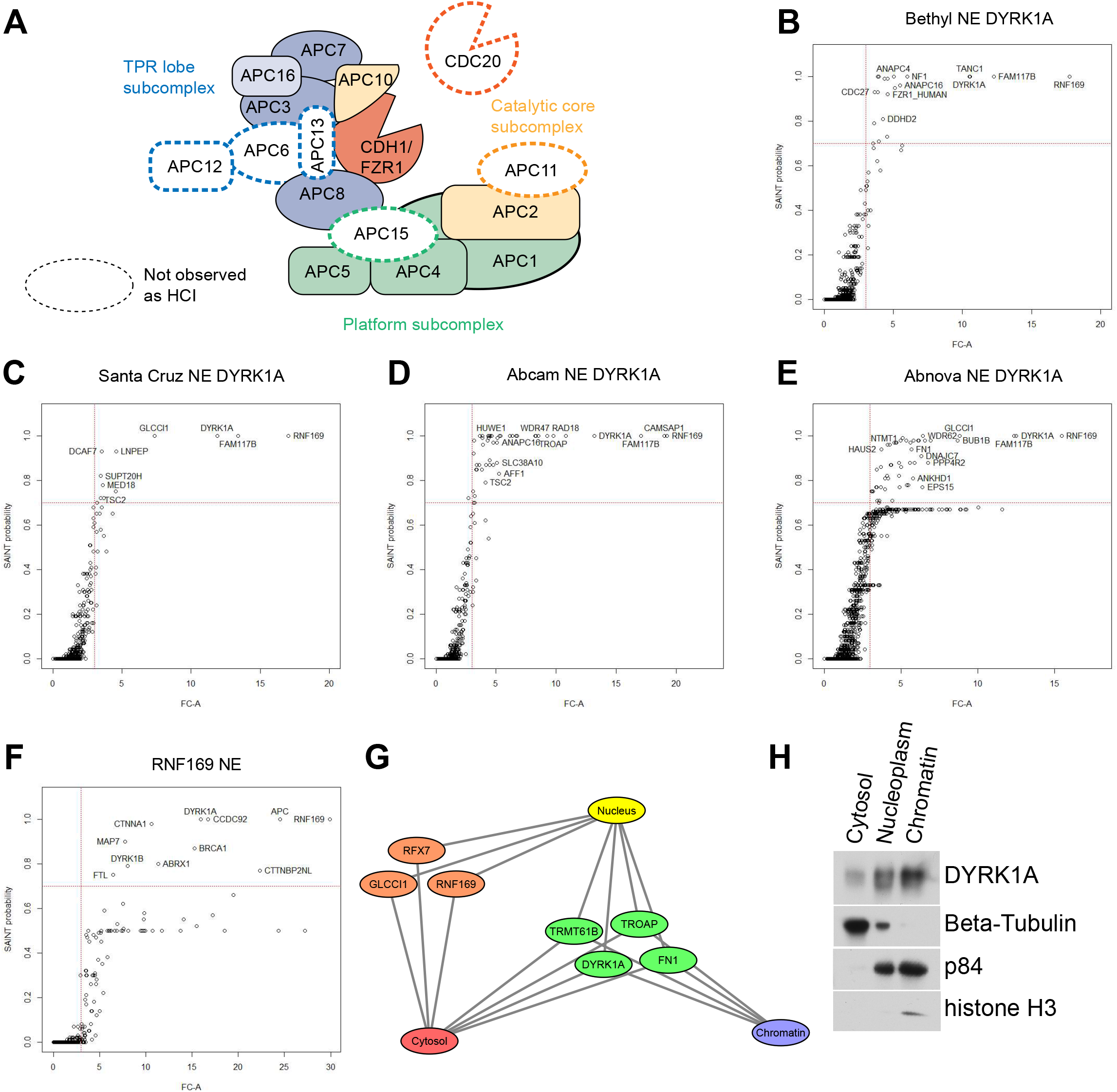

**Supplemental Figure 2.**
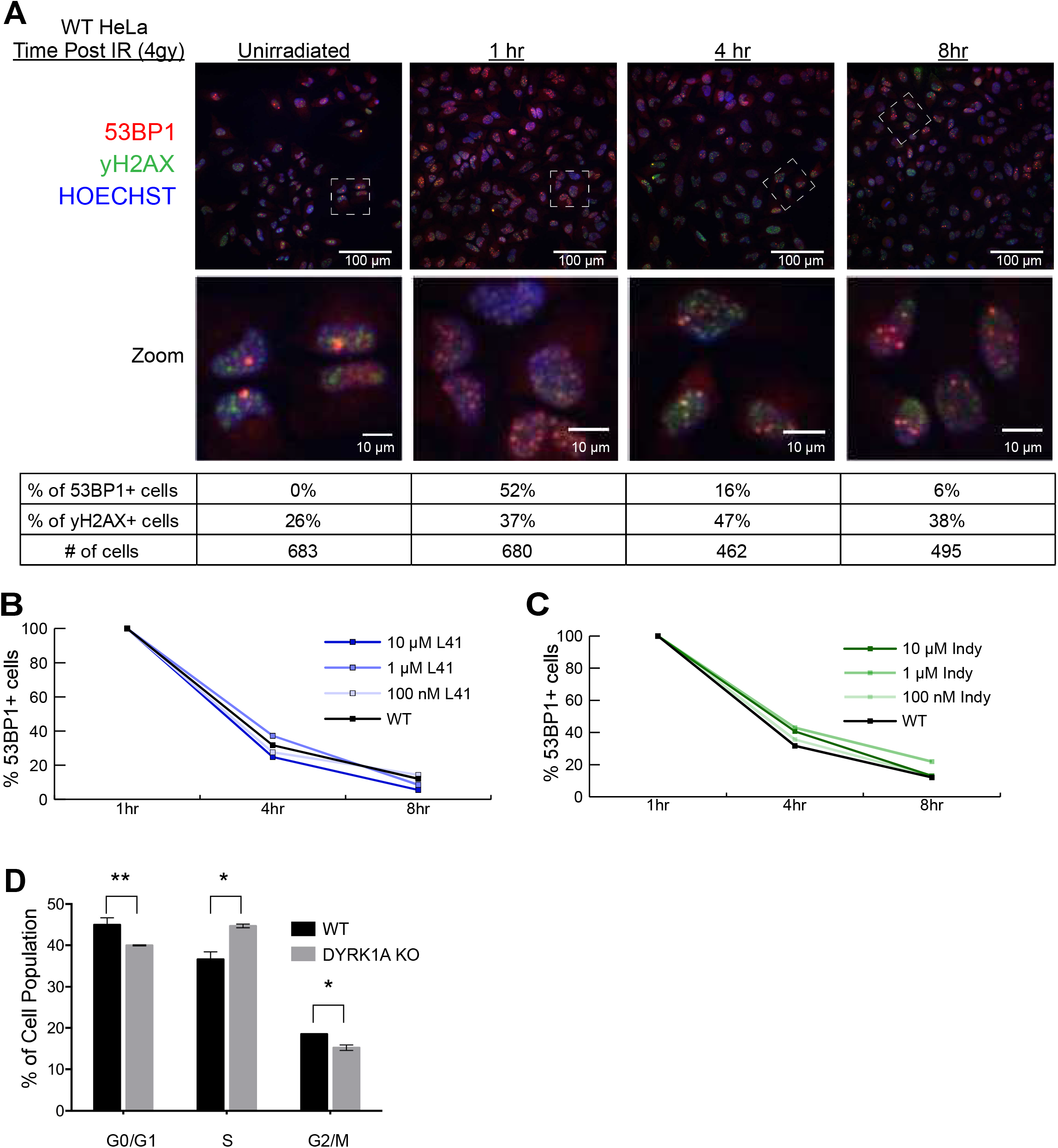

